# Febrile temperatures influence the transmission competence of mature *Plasmodium falciparum* gametocytes

**DOI:** 10.64898/2026.09.05.749077

**Authors:** Prince Chigozirim Ubiaru, Joyce Nyirongo, Antrea Pallikara, Lisa Ranford-Cartwright, Omar Janha

## Abstract

Malaria transmission depends on the survival and mosquito infectivity of mature *Plasmodium falciparum* gametocytes. Despite malaria febrile episodes reaching 39- 41.5°C in infected humans, the impact of febrile temperatures of varying durations on gametocyte viability and transmissibility remains undefined. Here we quantify the effects of exposing mature stage V gametocytes *in vitro* to febrile temperatures (39°C, 40°C, and 41.5°C) for varying durations (3-12 hours). To assess gametocyte morphology, functionality, and mosquito infectivity, we used light microscopy of Giemsa-stained thin blood smears, exflagellation assays, and standard membrane feeding assays (SMFAs) respectively.

Following exposure to 39°C, gametocytes retained normal morphology, exflagellation capacity, and mosquito infectivity across all exposure durations. At40°C, gametocytes remained morphologically intact and capable of exflagellation but exhibited a time-dependent reduction in mosquito infection, both in prevalence and oocyst intensity, following prolonged exposure. In contrast, exposure of mature gametocytes to 41.5°C, even for the minimum duration tested of 3 hours, resulted in complete loss of normal morphology, exflagellation capacity and ability to infect mosquitoes.

These findings reveal that mosquito infectivity of mature gametocytes is affected both by the magnitude of the febrile temperature and the duration of exposure. This provides insights into how host fever dynamics may influence parasite transmission.

## INTRODUCTION

Malaria remains a global public health issue, with *Plasmodium falciparum,* one of the five species infecting humans, responsible for most severe form of disease and deaths from malaria in sub-Saharan Africa. In 2024, the WHO reported approximately 282 million cases of malaria, resulting in an estimated 610, 000 deaths worldwide (World Health Organisation, 2025). Humans become infected when an infected female *Anopheles* mosquito takes a bloodmeal and injects sporozoites into the skin and bloodstream of the human host during feeding.

Fever, characterised by a rise in body temperature, rapid immune activation, increased inflammatory cytokines such as TNF-α (Tumour Necrosis Factor alpha), IFN-γ (Interferon gamma), and oxidative stress, is a common symptom of malaria. It is an important human host defence mechanism, triggered when schizont-infected erythrocytes rupture and release the merozoites, together with pyrogens such as haemozoin and glycosylphosphatidylinositol (Gazzinelli *et al.,* 2014; reviewed in Tint6-Font & Cortes, 2022).

During *P. falciparum* malaria, fever episodes are characterised by cyclical patterns known as paroxysms, typically occurring every 48 hours. During these episodes, body temperature rises from a normal baseline of about 37°C up to approximately 41°C and remains elevated for several hours. Within each paroxysm, temperatures often exceed 38.5°C for 2.5 to 10 hours. In some cases, brief spikes above 40°C may occur, usually lasting about 1 hour (Engelbrecht & Coetzer, 2013; reviewed in Tintó-Font & Cortes, 2022). However, in non-immune young children, fever episodes often present differently. Rather than following a regular cyclical pattern, children may experience sustained hyperpyrexia, frequently exceeding 40°C, which can persist for prolonged periods and vary in duration (reviewed in Schumacher & Spinelli, 2012). When temperature rises above an organism’s optimal range, proteins can misfold and aggregate, potentially resulting in cellular dysfunction, damage and death. Despite this challenge, malaria parasites can mount a heat shock response that helps maintain proteostasis under conditions exceeding their optimal growth temperature *(Rafols et al.,* 2026).

Febrile temperatures can be replicated *in vitro* by incubating asexual and gametocyte cultures at defined temperatures above 37°C. Different time intervals and febrile temperature ranges have been used to determine the effect of temperature on parasites, mimicking the temperature variation observed among different individuals during malaria fever (reviewed in Tintó-Font & Cortés, 2022). Most studies have observed that febrile temperatures damage or kill some of the intraerythrocytic stages of the parasite (Kwiatkowski, 1989; Long *et al.,* 2001), particularly the later stages of the asexual cycle. This often causes a decline in parasitaemia. However, the parasite ring stage often manages to survive. Overall, the effects of febrile temperatures on *P. falciparum* asexual blood stage parasites are well known. These include parasite growth inhibition, death, and synchronisation (Aunpad *et al.,* 2009; Engelbrecht & Coetzer, 2013; Gravenor & Kwiatkowski, 1998; Heidi *et al.,* 2008; Kwiatkowski, 1989; Kwiatkowski & Greenwood, 1989; Long *et al.,* 2001; Oakley *et al.,* 2007; Pavithra *et al.,* 2004; Singhaboot *et al.,* 2019; reviewed in Tintó-Font & Cortés, 2022). However, the effect of febrile temperature on mature gametocytes remains poorly understood.

Following sexual conversion, *P. falciparum* gametocytes take around 10-12 days to reach maturity (Neveu *et al.,* 2018; Rogers *et al.,* 2000), during which period the first four stages are sequestered in the deep tissues such as the bone marrow and spleen (Aguilar *et al.,* 2014; Joice *et al.,* 2014; Rogers *et al.,* 2000; Smalley, Abdalla, & Brown, 1981), only emerging to circulate in the blood as mature stage V gametocytes. The exact period in which mature stage V gametocytes circulate in the blood after release from the bone marrow is not known but has been estimated by mathematical modelling to be a median of 6 days (Bousema *et al.,* 2010; Eichner *et al.,* 2001). This suggests that mature gametocytes are likely to experience one febrile temperature episode during peripheral circulation in symptomatic individuals.

Field studies have shown that gametocyte carriers with fever at the time of transmission are less infectious to mosquitoes than afebrile carriers (Ahmad *et al.,* 2021; Barry *et al.,* 2021; Gouagna *et al.,* 2004). Mosquito infectivity is influenced by multiple mammalian host and parasite factors such as mature gametocyte density, transmission-blocking immunity, inflammatory cytokines, anaemia and parasite genotype, making it difficult to attribute fever to the observed reduction in transmission with field studies. Nonetheless, experimental evidence supports a direct negative effect of elevated temperature on transmission. For instance, incubating mature *P. falciparum* gametocytes at 42°C for either 15 minutes or 4 hours has been shown to reduce mosquito infection rates (Soumare *et al.,* 2021). Furthermore, stage V gametocytes exposed to 41°C for one hour were reported to lack the ability to exflagellate normally (Ràfols *et al.,* 2026), and the authors suggested that patients experiencing severe fever may therefore become temporarily non-infectious to mosquitoes.

Although it is established that young asexual parasites in early developmental stages (young rings) possess fundamental biological properties that help them propagate successfully within the human host under febrile conditions, these conditions are known to inhibit the growth of later stages of the parasite such as trophozoites and schizonts but increase sexual conversion and gametocytaemia of the parasites *in vitro* (Portugaliza *et al.,* 2020). However, it remains unknown whether mature gametocytes can withstand similar febrile conditions. No study has established the febrile temperature or heat shock threshold, the duration of exposure required, or the conditions under which mature gametocytes lose their ability to infect mosquitoes. This is partly due to the prevailing assumption that gametocyte transmissibility is compromised at febrile temperatures, as well as the technical challenges associated with mosquito infection assays. This lack of evidence represents a critical gap in our understanding of malaria transmission biology, given that only mature male and female gametocytes can infect mosquito vectors.

To address this gap, we investigated the febrile temperature thresholds and exposure duration that compromise the ability of mature gametocytes to infect mosquitoes. Stage V gametocytes were exposed to a range of temperatures representative of febrile conditions across different time intervals.

## MATERIALS AND METHODS

### Malaria parasite culture

Parasites were maintained in human blood (blood group 0), at 5% haematocrit in complete RPMI with 0.5% w/v albumax and 1x ITS-X (Pradhan, Ubiaru, & Ranford-Cartwright, 2024), at 37°C, under a gas mixture of 96% nitrogen, 3% carbon dioxide and 1% oxygen. Gametocyte cultures were set up according to standard protocols with slight modifications (Carter, Ranford-Cartwright, & Alano, 1993; Pradhan, Ubiaru, & Ranford-Cartwright, 2024). For each feed and temperature condition, two 5 ml cultures at 6% haematocrit, 0.5% parasitaemia were set up in T-25 flasks (Corning, Cat No-430372) and maintained for days 15 and days 17 respectively. The media was increased to 7.5 ml (“bulking-up”) on day 3, once the parasites were visibly stressed, a sign of sexual commitment. From day 7, the cultures were treated for 3 days with 20 U/ml heparin (Merck H3149) dissolved in the medium, to inhibit erythrocyte reinvasion and thereby prevent further development of the asexual stages (Miao et al., 2013). The media in the culture flasks were changed daily without disturbing the settled erythrocytes at the bottom of the flask until they were used for infectious feeds as a mixture of cultures at 15 and 17 days after setup.

## EXPOSURE TO DIFFERENT TEMPERATURES/ DURATIONS

To test the effect of febrile temperatures on the ability of gametocytes to remain viable, functional and infectious to mosquitoes, gametocytes were exposed to various elevated temperatures. The duration of the exposure to higher temperatures varied independently for each temperature condition tested. On day 14 (D14), separate gametocyte cultures were incubated at 39°C, 40°C and 41.5°C for 3, 6, and 12 hours in individual incubators, while control cultures were maintained at 37°C (Figure 1). These exposure times were selected because it has been established that fever episodes of >38.5°C up to >40°C can last for between 1 and 10 hours (Tintó-Font & Cortes, 2022). Following exposure to febrile temperatures the cultures were returned to standard incubation condition (37°C) and maintained until the day of the infectious feed.

**Figure 1.**
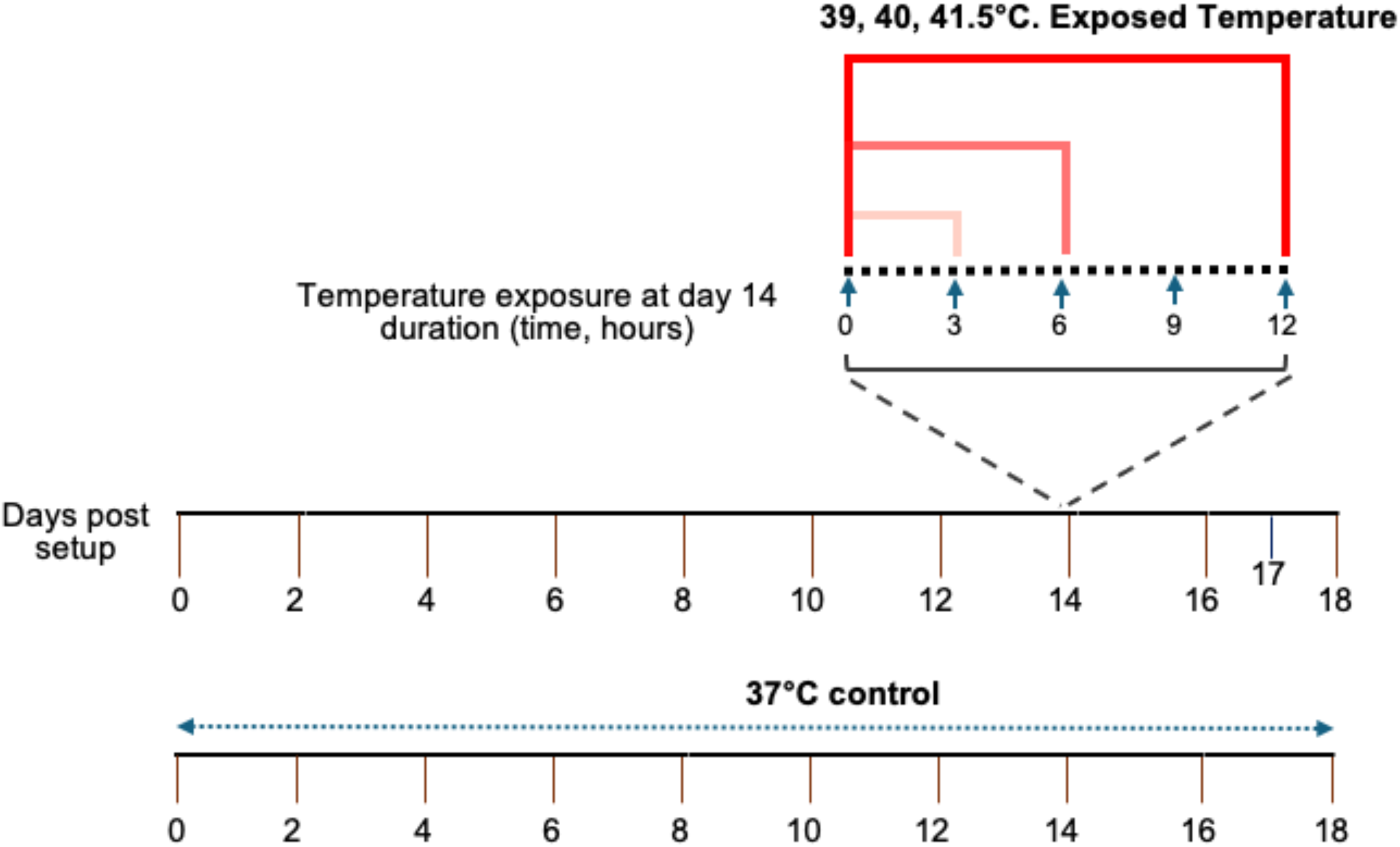
Experimental design. Schematic representation of the experimental design showing the exposure windows for *P. falciparum* gametocytes at febrile temperatures of 39°C, 40°C, and 41.5°C. On day 14, gametocytes were exposed to each temperature for 3, 6, or 12 hand maintained at 37°C under standard culture conditions outside the exposure periods. Control gametocytes were maintained continuously at 37°c.

### Mosquito rearing

Female mosquitoes aged 5-6 days post-emergence were used for infectious feeds with the treated gametocyte cultures. Adult female *Anopheles coluzzii* (N’Gousso line) mosquitoes were reared under controlled insectary conditions at 26-28°C, 70-80% relative humidity, and a 12-hour light: 12-hour dark photoperiod. Larvae were maintained and fed daily with a finely ground larval diet. Adults were collected and transferred to mosquito cups and were maintained on a 5% glucose solution+ 0.05% w/v 4-amino benzoic acid continuously. A night before the feeding, the mosquitoes were starved (glucose was removed and replaced with water) to enhance their feeding.

### Standard membrane feeding assays (SMFAs)

*An. coluzzii* (N’Gousso) mosquitoes were offered a mixture of day-15 and day-17 old gametocytes via membrane feeder according to standard protocols (Carter, Ranford-Cartwright, & Alano, 1993), with the final gametocytaemia in the blood meals adjusted to be in the same range of 0.6-1% gametocytaemia where the gametocytes were present. For each elevated temperature condition, four cups of mosquitoes were allowed to feed on the gametocytes, one each for parasites that had been exposed for 3, 6 and 12h and one for the control that had been maintained at 37°C. The experiment was repeated twice for each temperature/exposure time condition.

Blood-fed mosquitoes were maintained at 26-28°C, 70-80% relative humidity and were given fresh glucose (5%) containing 0.05% w/v 4-amino benzoic acid *ad libitum.* Mosquitoes were dissected 9-10 days post blood meal to visualise and count oocysts on the midguts (without staining) under 400X magnification. Infection prevalence (percentage of mosquitoes infected) and oocyst intensity (number of oocysts per midgut) were recorded. A total of 28-32 mosquitoes were examined for each mosquito cup.

### Preparation of gametocyte smears

Thin smears were prepared from the infectious feed to determine the morphology and number of gametocytes in the blood meal by microscopy. The smears were air-dried, fixed with absolute methanol, and stained with 20% Giemsa stain for 15 min.

### Exflagellation assay

A drop of the infectious bloodmeal with the temperature-exposed gametocyte culture was placed on a glass slide at room temperature to allow gamete activation, covered with a coverslip, and observed under 400X magnification after 15-30 minutes at room temperature. The presence or absence of exflagellation centres characterised by actively motile microgametes radiating from a central body was observed for each temperature condition.

### Morphological analysis of mature gametocytes

Thin blood smears were examined under light microscopy at 1000X magnification, and gametocytes were counted by examining at least 1000 erythrocytes per blood film. Each mature stage V gametocyte was classified as morphologically normal (if it retained the characteristic elongated crescent shape with intact membrane and visible pigment granules) or morphologically abnormal (if it exhibited cellular shrinkage, loss of crescent shape, membrane irregularity, or rounding).

### Statistical analysis

The impact of exposure to elevated temperatures on the number of gametocytes and mosquito infection prevalence was analysed with generalised linear models (GLMs) using R (version 4.3.0) (Ubiaru & Ranford-Cartwright, 2025). Gametocyte number was analysed using a GLM with a Poisson distribution and log link, with exposure temperature, exposure time (h) and replicate included as fixed effects. Mosquito infection prevalence was analysed using a binomial GLM with a logistic regression, with exposure temperature, exposure time (h), replicate and gametocyte number in the infectious bloodmeal (gametocytes per 1000 RBC) included as fixed effects. Non-significant variables were removed by backward elimination (Burnham, 1998), and the final minimal statistically significant model was used to obtain the estimated number of gametocytes and infection prevalence with 95% confidence intervals, and the significance of the difference in number of gametocytes and infection prevalence between different temperatures and exposure time durations.

Intensity of infection (oocyst numbers per mosquito) was analysed with negative binomial (NB), zero-inflated (ZINB) and hurdle negative binomial (Hurdle NB) regression models using the R package pscl (Jackman, 2024). This was done to account for the overdispersion and excess zeroes commonly seen in oocyst intensity data (Ubiaru & Ranford-Cartwright, 2025). The maximal model contained the fixed variables of exposure temperature, time (h), replicate and gametocyte numbers (gametocytes per 1000 rbc). Again, non-significant variables were removed by backward elimination. The best fit model from (NB, ZINB and Hurdle NB) was determined by comparison of AIC and the accuracy of prediction of the number of zero values (Ubiaru & Ranford-Cartwright, 2025). The final minimal statistically significant and best fit model (ZINB) was used to obtain the estimated oocyst intensity and 95% confidence intervals, and the significance of the difference in intensity between different exposure temperatures and exposure time durations.

## RESULTS

### Impact of exposure to elevated temperature on gametocyte morphology

To determine if exposure to elevated temperatures had an adverse effect on mature Stage V gametocytes, gametocyte density and morphology were assessed using Giemsa-stained thin blood smears. No morphological abnormalities were seen in gametocytes exposed to temperatures of 39°C and 40°C, for either 3, 6 or 12 hours, compared to gametocytes not exposed to these elevated temperatures (Figures 2 & 3 a-i-iv). There was no statistically significant difference in gametocyte number following exposure to elevated temperatures of 39°C or 40°C for 3, 6, or 12 hours and those maintained at 37°C (Supplementary table 1a & b). All stage V gametocytes retained the characteristic elongated crescent morphology typically associated with normal mature stage V parasites before and after temperature exposure at 39°C and 40°C (Figures 2 & 3 a-i-iv). However, exposure to a temperature of 41.5°C resulted in altered morphology such as cellular shrinkage and loss of the characteristic crescent shape under microscopic examination (Figure 4a-ii-iv). There was a significant decrease in gametocyte number between gametocytes exposed to temperature of 41.5°C and those maintained at 37°C (Supplementary table 1c).

**Figure 2.**
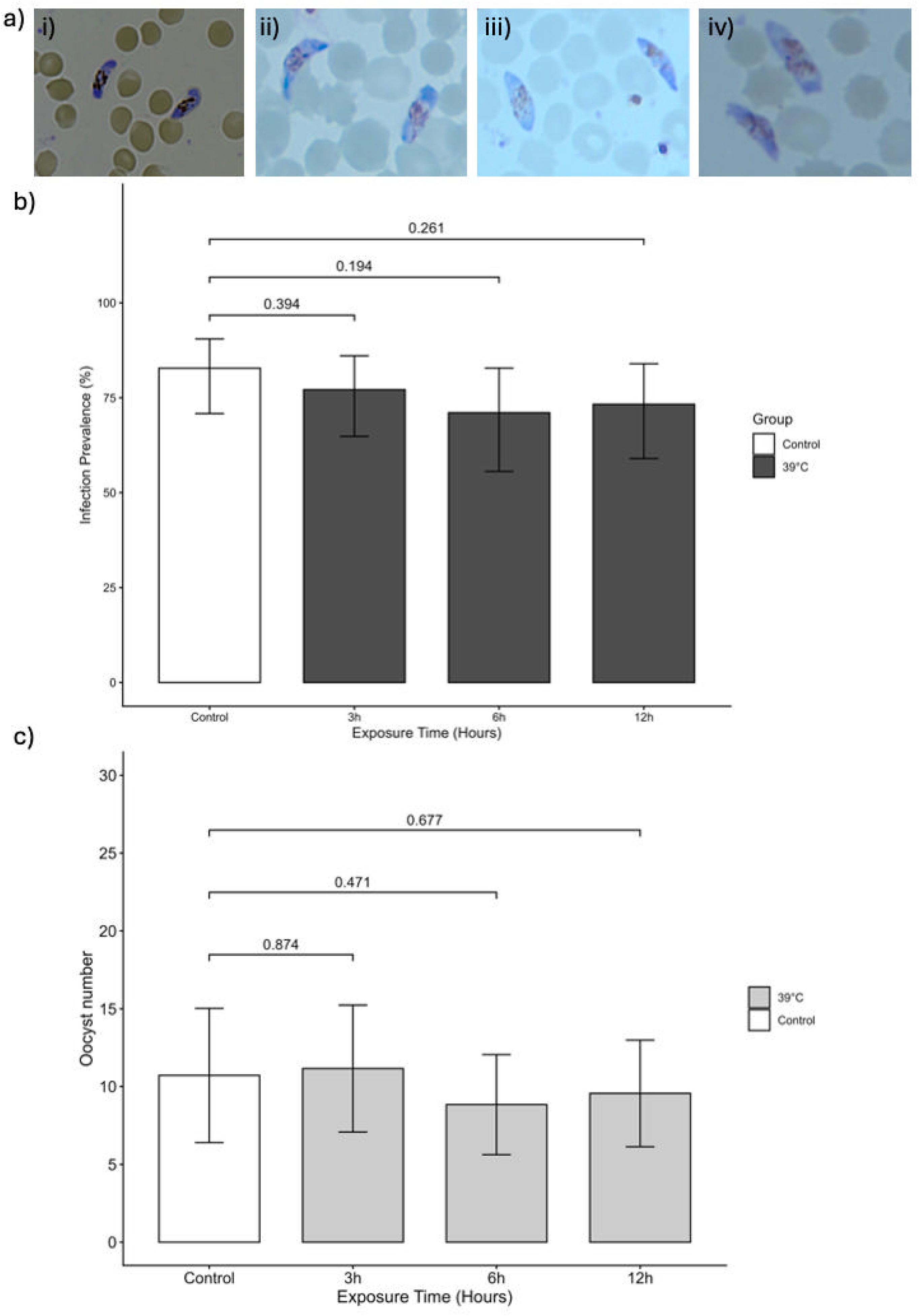
Effects of mild fever on mature gametocytes and mosquito infectivity. (a) Representative images of stage V gametocytes maintained at 37°C (control; a-i) or exposed to 39°C for 3, 6, and 12 h (a-ii-iv), showing normal, elongated crescent-shaped morphology without visible defects. Scale bar, 20 µm. (b) Mosquito infection prevalence and (c) oocyst intensity following exposure of stage V gametocytes to 39°C for 3, 6, and 12 h. Infection prevalence was estimated using a generalized linear model (GLM), and oocyst intensity using a zero-inflated model. *P* values were obtained from the respective models; error bars represent 95% confidence intervals. *n* = 2 independent replicates per exposure duration.

**Figure 3.**
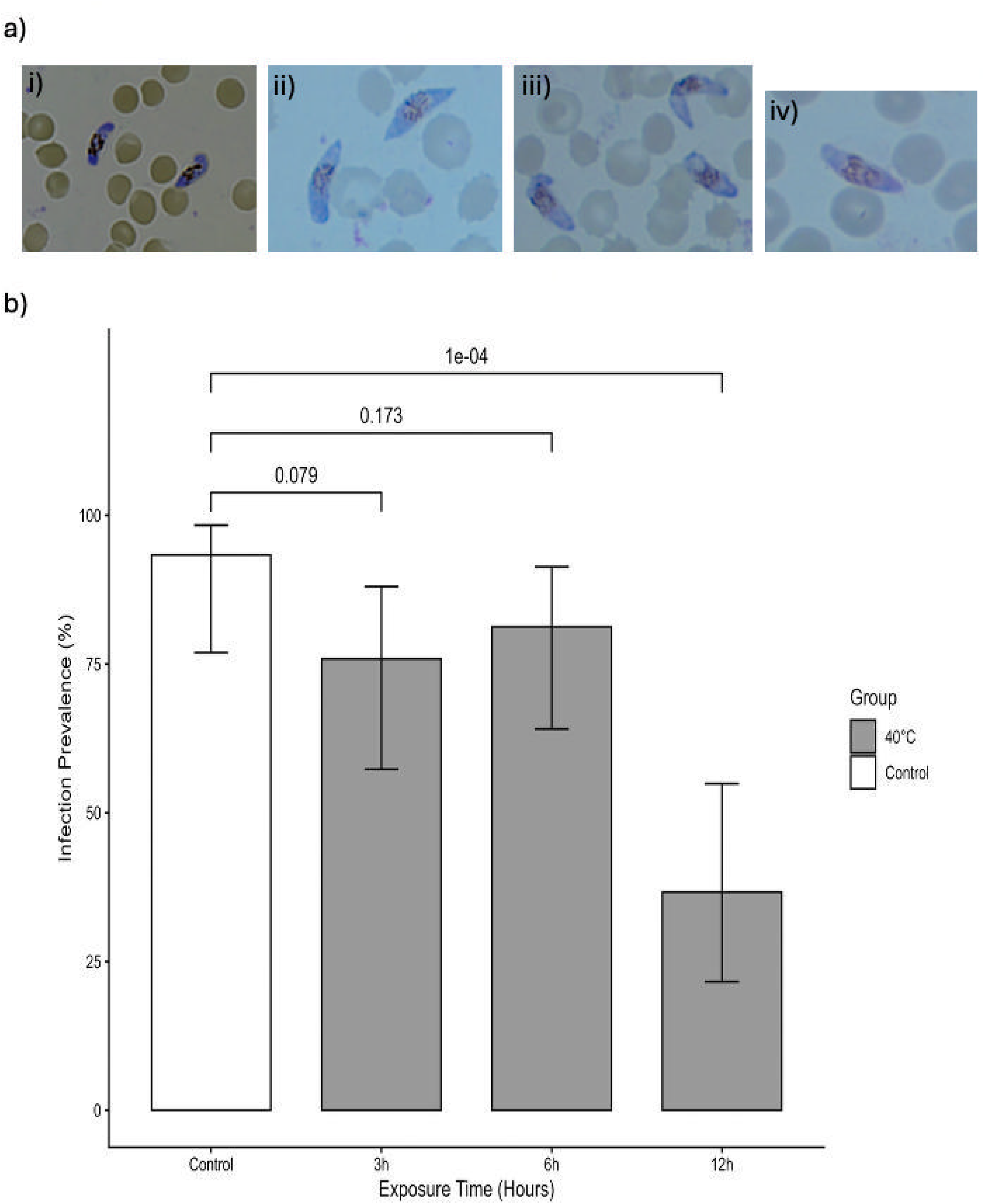
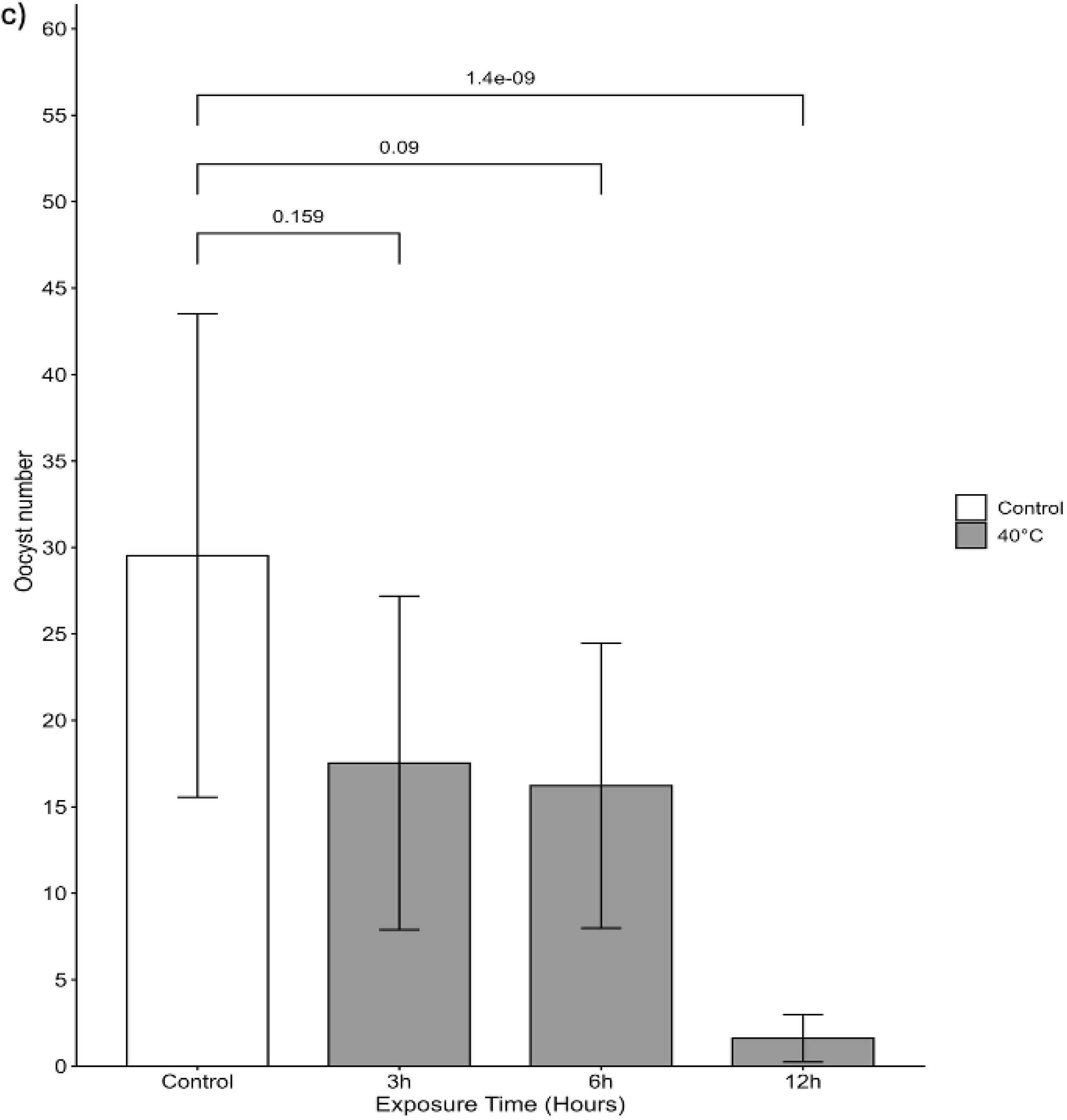
Effects of mild fever on mature gametocytes and mosquito infectivity. (a) Representative images of stage V gametocytes maintained at 37°C (control; a-i) or exposed to 40°C for 3, 6, and 12 h (a-ii-iv), showing normal crescent-shaped morphology without visible defects. Scale bar, 20 µm. (b) Mosquito infection prevalence and (c) oocyst intensity following exposure of stage V gametocytes to 40°C for *3, 6,* and 12 h. Infection prevalence was estimated using a generalized linear model (GLM), and oocyst intensity usinga zero-inflated model. *P* values were obtained from the respective models; error bars represent 95%>confidence intervals. *n* = 2 independent replicates per exposure duration.

**Figure 4.**
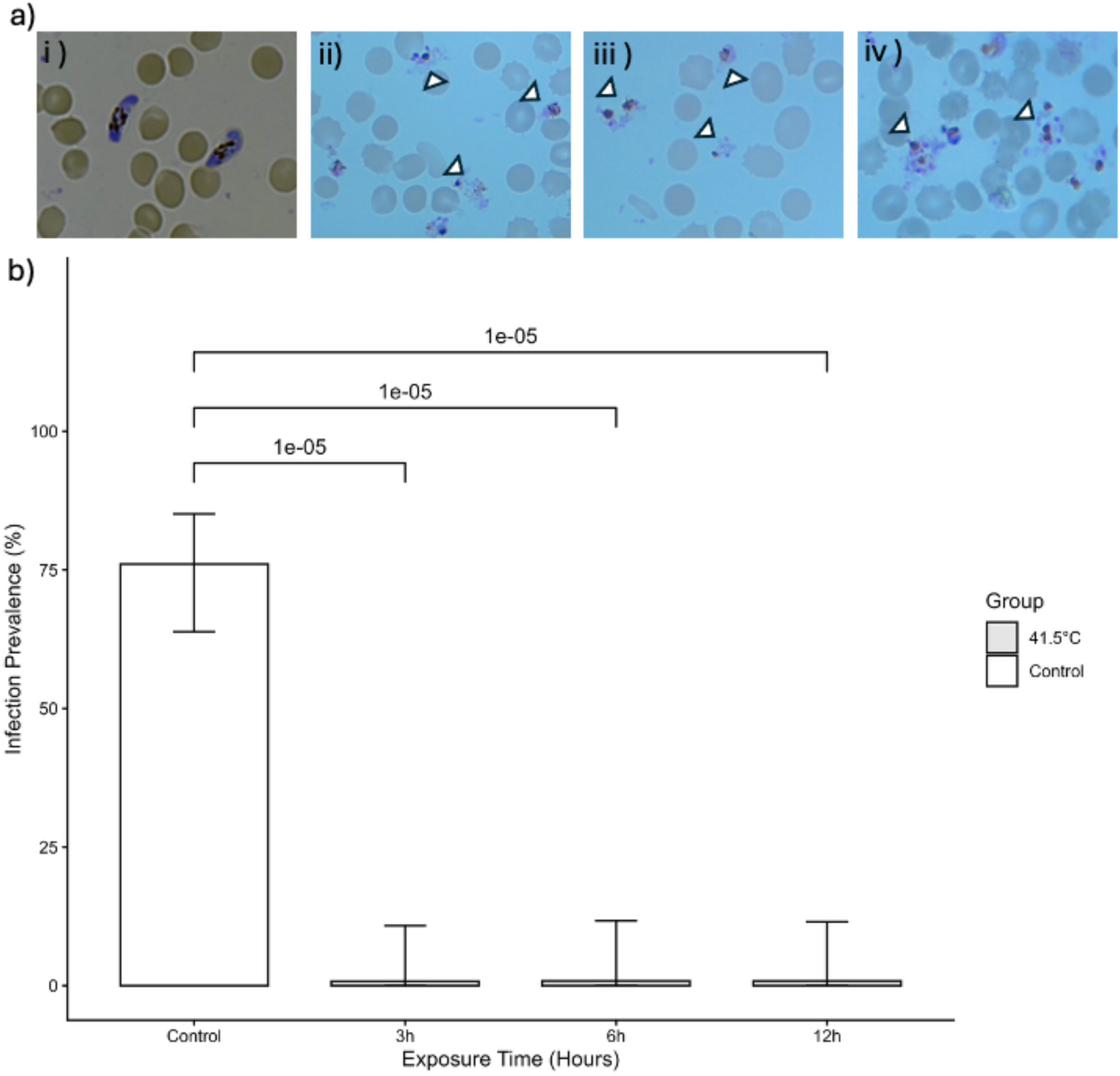
Effects of severe fever on mature gametocytes and mosquito infectivity. **(a)** Representative images of stage V gametocytes maintained at 37°C (control; a-i) or exposed to 41.5°C for *3,* 6, and 12 h (a-ii-iv). Control gametocytes show normal elongated, crescent-shaped morphology. Heat-exposed gametocytes show substantial lysis and morphological abnormalities (white triangular arrows). Scale bar, 20 µm. **(b)** Mosquito infection prevalence after exposure of stage V gametocytes to 41.5°C for 3, 6, and 12 h. Infection prevalence was estimated using a generalized linear model (GLM) with logistic regression fitted using *logistf. P* values were obtained from the respective models; error bars represent 95% confidence intervals. *n* = 2 independent replicates per exposure duration.

This observed phenotype is consistent with the effects of high temperature on living cells due to protein denaturation and activation of heat shock proteins that would trigger necrosis (Bellmann et al., 2010; Hu et al., 2022; Kayastha et al., 2024).

### Impact of exposure to elevated temperature on exflagellation

To assess whether exposure to elevated temperatures affects the exflagellation capacity of mature stage V male gametocytes, exflagellation was checked using an exflagellation assay. In the blood meal preparation from cultures not exposed to elevated temperatures, exflagellation centres were seen within 12 minutes of making the slide, and continued for ∼10 minutes, with 1 to 5 exflagellation events per field of view under 4000x magnification. The same timing and level of exflagellation events were seen for gametocytes exposed to 39°C or 40°C for 3, 6 or 12 hours. In contrast, no exflagellation events were observed when the cultures had been exposed to 41.5°C for 3, 6 or 12 hours, even when the slide was examined for 20-30 minutes.

### Impact of exposure to elevated temperature on mosquito infectivity

Mosquito infection prevalence was similar between the control and gametocytes exposed to elevated temperature of 39oC (Figure 2b). There was no significant difference (P>0.05) in mosquito infection prevalence with gametocytes exposed to 39°C for 3h, 6h or 12h, compared with gametocytes maintained at 37°C. For oocyst intensity, none of the oocyst infection levels derived from gametocytes exposed to temperatures of 39°C differed significantly from the 37°C control (P>0.05 for all, Figure 2c).

Gametocytes exposed to a temperature of 40°C for 12h gave a significantly lower prevalence of infection compared to those maintained at 37°C (P=0.0001), but a shorter temperature exposure of 3h or 6h did not impact significantly on infection prevalence (P= 0.079 (3h), P=0.173 (6h) Figure 3b). Similarly, oocyst intensity in infections with gametocytes exposed to a temperature of 40°C for 12h was significantly lower compared to gametocytes maintained at 37 °C (P=1.4 x 1o-^9^), but a shorter temperature exposure of 3h or 6h did not impact significantly on oocyst intensity (P= 0.159 (3h), P=0.09 (6h); Figure 3c).

Gametocytes exposed to a temperature of 41.5°C for all the time durations (3h, 6h and 12h) resulted in a highly significant reduction in infection prevalence to zero (P = 0.00001 (3h, 6h, 12h) compared to gametocytes maintained at 37°C (Figure 4b). As there was no infection in the mosquitoes fed gametocytes exposed to 41.5°C, it was not possible to analyse infection intensity.

## DISCUSSION

The results presented here show that mature *P. falciparum* stage V gametocytes can survive moderate febrile temperatures for a short period and still retain their ability to infect mosquitoes. However, the parasites progressively lose their ability to infect mosquitoes as temperatures increase, and with longer exposure times.

Exposure of mature gametocytes to 39°C for up to 12h had no impact on gametocyte morphology, their ability to exflagellate, or on mosquito infectivity, whereas infection was completely prevented after exposure of gametocytes to 41.5°C, even for the minimum duration of 3h, with significant impacts on morphology and exflagellation.

At 40°C exposure, neither gametocyte morphology nor exflagellation was affected. However, the exflagellation assessment was qualitative and therefore may not fully capture the impact of temperature exposure on male gametocyte fitness. Although exflagellation was observed following exposure to 40°C for up to 12h, mosquito infectivity and oocyst intensity were significantly reduced after 12h, but not after shorter exposure durations of 3h or 6h. This discrepancy suggests that qualitative assessment of exflagellation may underestimate temperature-induced effects on transmission competence. Alternatively, prolonged exposure to 40°C may have a greater impact on female gametocyte fitness, which cannot be assessed directly. As successful mosquito infection depends on the functional competence of both male and female gametocytes, impairment of either sex could reduce transmission despite apparently normal male exflagellation. These findings suggest that exposure to moderate or mild febrile temperatures for short time periods (6h or less) does not affect the survival of mature gametocytes or their ability to infect mosquitoes, whereas longer exposures to these moderately elevated temperatures did affect gametocyte infectivity. This is consistent with previous reports showing that short-term exposure (only 15 minutes) of gametocytes to 40°C did not affect mosquito infection (Soumare *et al.,* 2021), and stage V gametocytes survive a 1-hour heat shock at 41°C without activating a heat shock response (Ràfols *et al.,* 2026). Together, these findings suggest that tolerance of elevated temperatures could be an important adaptive trait of mature gametocytes, which must persist in the human bloodstream for some time despite recurrent febrile episodes during malaria infection.

Exposure to 41.5°C even for short periods of time (3h) produced a rapid loss of gametocyte viability. Stage V gametocytes displayed morphological degeneration; no exflagellation was observed, and mosquito infections were completely ablated across all exposure durations, including as short as only 3 hours exposure to the elevated temperature. Other researchers have recently reported a detrimental impact of a short exposure (1h) to a 41°C temperature on male gametocyte exflagellation, although they reported that stage V gametocytes looked morphologically normal (Ràfols *et al.,* 2026). Similarly, a 15-minute incubation of a gametocyte-containing blood meal at 42°C was reported to inactivate gametocytes, leading to a near complete blockade of mosquito transmission (Soumare *et al.,* 2021). Our findings indicate that exposure to temperatures in the hyperpyrexic range (>41°C) exceed the temperature tolerance of mature stage V gametocytes and can rapidly eliminate their infectivity (Figure 4b). At high grade fever temperatures of ≥40°C, a longer exposure time of 12h is required to significantly decrease infection rates in mosquitoes (Figure 3b & 3c), and exposure for up to 12h at moderate fever levels of 39°C had no effect on gametocyte infectivity (Figure 2b & 2c). This transition from no impact, through partial impairment at 40°C and complete loss of infectivity at 41.5°C suggests a temperature limit, beyond which mature gametocytes can no longer successfully infect mosquitoes.

Altogether, these findings have important implications for malaria transmission biology. Firstly, febrile temperatures up to 39°C are unlikely to substantially reduce transmission from individuals carrying mature gametocytes. Previous studies in natural infections have reported lower gametocyte infectivity among gametocyte carriers with fever (> 37.5°C) at the time of mosquito feeding (Ahmad *et al.,* 2021; Barry *et al.,* 2021; Gouagna *et al.,* 2004). Second, sustained higher febrile temperatures of 40°C may partially or totally suppress transmission, particularly when the fever duration is prolonged. Thirdly, hyperpyrexic temperatures (above 41.5°C) are predicted to profoundly reduce gametocytes’ infectiousness and block transmission. Therefore, human host febrile temperature may influence transmission, with modest fever exerting little effect and severe hyperthermia acting as a natural transmission-limiting factor.

These results may also help explain the variability in gametocyte infectivity observed in naturally infected individuals with fever (Ahmad *et al.,* 2021; Barry *et al.,* 2021; Gouagna *et al.,* 2004), as differences in fever intensity, duration, timing relative to gametocyte maturity, and use of antipyretic treatment could all alter the transmission potential of circulating gametocytes. In endemic settings, suppression of fever through early treatment may improve clinical outcomes but could theoretically preserve gametocyte infectivity if temperature otherwise reduces transmission.

Some limitations should be acknowledged. The study was conducted under controlled *in vitro* conditions using defined temperatures and exposure durations, which may not fully replicate fluctuating *in vivo* fever patterns in human host.

Future work could investigate molecular mechanisms underlying temperature tolerance and injury. It would also be valuable to assess whether repeated febrile episodes of shorter duration cumulatively impair gametocyte infectivity and whether different parasite clones differ in temperature resilience. Understanding these processes may clarify how host fever shapes malaria transmission under natural conditions.

## CONCLUSIONS

Mature stage VP. *falciparum* gametocytes exhibit marked tolerance to moderate fever range temperatures, remaining viable and fully infectious following exposure to 39°C for up to 12 hours. In contrast, prolonged exposure to 40°C significantly compromises transmission success, whereas exposure to 41.5°C rapidly affects both gametocyte viability and mosquito infectivity. These findings reveal a graded temperature response in mature gametocytes, in which 39°C is well tolerated, 40°C induces time-dependent reductions in infectivity, and 41.5°C eliminates transmission competence. Our results highlight host febrile temperature intensity and duration may modulate malaria transmission potential in a temperature dependent manner.

## DATA AVAILABILITY

Data are available from the corresponding authors upon request.

## ACKNOWLEDGEMENTS

We want to thank Susanne Krabbendam for assistance with maintaining the SBOHVM Glasgow University mosquito insectaries; the Scottish National Blood Transfusion Service for providing human blood; and Dr Sabyasachi Pradhan for his input during the conception of this research and for comments on the draft manuscript. We also acknowledge and thank Anne Lawson for assisting in transferring the gametocyte culture flasks from the experimental incubator to the control incubator.

## AUTHOR CONTRIBUTIONS

PU conceived the idea, performed all the experiments, conducted the statistical data analysis, and wrote the first draft. OJ conceived the idea, designed the experiments, performed all the experiments, wrote the first draft, supervised and coordinated the team, and reviewed the final versions of the figures and manuscript prior to submission. LRC advised on the study concept and experimental design, reviewed the manuscript, and provided revisions that substantially improved the final manuscript. **JN** acquired the microscopy images used in the manuscript and assisted with the experiments. AP assisted with the laboratory experiments. All authors read and approved the final manuscript.

## FINANCIAL SUPPORT

This work did not receive any specific funding. PU was supported by a British Council International Science Partnerships Fund (ISPF) grant awarded to Lisa Ranford-Cartwright, while Omar Janha was supported by funding from the Gates Foundation (INV-039928 and INV-088781).

## COMPETING INTERESTS

The authors declare no competing interests.

## ETHICAL STANDARDS

The use of human blood and serum was approved by the Scottish National Blood Transfusion Service Committee for the Governance of blood and tissue samples for non- therapeutic use (Sample Submission Reference No. 23 ∼ 11) and by the University of Glasgow College Ethics committee (Project No: 200220442).

## Notes

### Competing Interest Statement

The authors have declared no competing interest.

