## Supplementary Table for "Febrile temperatures influence the transmission competence of mature *Plasmodium falciparum* gametocytes"

Table 1a. GLM analysis of differences in gametocyte numbers between the 37°C control and 39°C treatments (3, 6, and 12 h)

| Temperature & time duration (hours) | Gametocyte Number (G'C per 1000 RBC) | | GLM Predicted Gametocyte number | P value |
|---|---|---|---|---|
| | Replicate 1 | Replicate 2 | | |
| 37_control | 6.0 | 10 | 8.0 | |
| 39_3 | 6.0 | 9.7 | 7.9 | 0.947 |
| 39_6 | 6.0 | 7.6 | 6.8 | 0.603 |
| 39_12 | 6.0 | 8.0 | 7.0 | 0.663 |

Table 1b. GLM analysis of differences in gametocyte numbers between the 37°C control and 40°C treatments (3, 6, and 12 h)

| Temperature & time duration (hours) | Gametocyte Number (G'C per 1000 RBC) | | GLM Predicted Gametocyte number | P value |
|---|---|---|---|---|
| | Replicate 1 | Replicate 2 | | |
| 37_control | 6.0 | 9.0 | 7.5 | |
| 40_3 | 9.0 | 7.0 | 8.0 | 0.733 |
| 40_6 | 7.0 | 8.0 | 7.5 | 1.000 |
| 40_12 | 6.0 | 7.0 | 6.5 | 0.505 |

Table 1c. GLM analysis of differences in gametocyte numbers between the 37°C control and 41.5°C treatments (3, 6, and 12 h)

| Temperature & time duration (hours) | Gametocyte Number (G'C per 1000 RBC) | | GLM Predicted Gametocyte number | P value |
|---|---|---|---|---|
| | Replicate 1 | Replicate 2 | | |
| 37_control | 6.0 | 10 | 8.0 | |
| 41.5_3 | 0 | 0 | 0 | **0.005 **** |
| 41.5_6 | 0 | 0 | 0 | **0.005 **** |
| 41.5_12 | 0 | 0 | 0 | **0.005 **** |
